# The protein phosphatase-2A subunit PR130 is linked to cytotoxic protein aggregate formation in mesenchymal pancreatic ductal adenocarcinoma cells

**DOI:** 10.1101/2023.09.03.556106

**Authors:** Alexandra Nguyen, Alessa K. Leydecker, Al-Hassan M. Mustafa, Janine Murr, Falk Butter, Oliver H. Krämer

**Affiliations:** Department of Toxicology, University Medical Center, Obere Zahlbacher St. 67, 55131 Mainz, Germany; Department of Zoology, Faculty of Science, Aswan University, Aswan, Egypt; Medical Clinic and Polyclinic II, Klinikum rechts der Isar, Technical University Munich, 81675 München, Germany; Institute of Molecular Biology (IMB), Ackermannweg 4, 55128 Mainz, Germany; Federal Research Institute for Animal Health, Südufer 10, 17493 Greifswald - Insel Riems

## Abstract

Protein phosphatase-2A (PP2A) is a major source of cellular serine/threonine phosphatase activity. PP2A B-type subunits regulate the intracellular localization and the catalytic activity of PP2A-A/PP2A-C complexes towards individual proteins. There is limited knowledge on how PP2A B-type subunits regulate biologically important functions and if these subunits determine the growth and drug responsiveness of tumor cells. Pancreatic ductal adenocarcinoma (PDAC) is a dismal disease with poor prognosis. Mesenchymal PDAC subtypes are more aggressive and metastasis-prone than epithelial subtypes. We show that mesenchymal PDAC cells express significantly higher levels of the PP2A B-type subunit PR130 and its mRNA *Ppp2r3a* than epithelial PDAC cells (n=38). Among 17 PP2A B-type subunits, this differential regulation is unique for *Ppp2r3a* and PR130. The higher levels of PR130 in mesenchymal PDAC cells are linked to their vulnerability to the PP2A inhibitor phendione. Phendione induces apoptosis and an accumulation of cytotoxic protein aggregates in such cells. These processes occur independently of the major tumor suppressor p53, which is frequently mutated in PDAC cells. Proteomic analyses reveal that phendione upregulates the chaperone heat shock protein HSP70 in mesenchymal PDAC cells. Inhibition of HSP70 promotes phendione-induced apoptosis. We additionally disclose that phendione promotes a proteasomal degradation of PR130. Genetic elimination of PR130 sensitizes mesenchymal PDAC cells to phendione-induced apoptosis and protein aggregate formation. These data illustrate pharmacologically amenable, selective dependencies of mesenchymal PDAC cells on PP2A-PR130 and HSP70. PP2A inhibition triggers a harmful accumulation of protein aggregates in neurons. This undesired mechanism might be exploited to kill mesenchymal tumor cells.

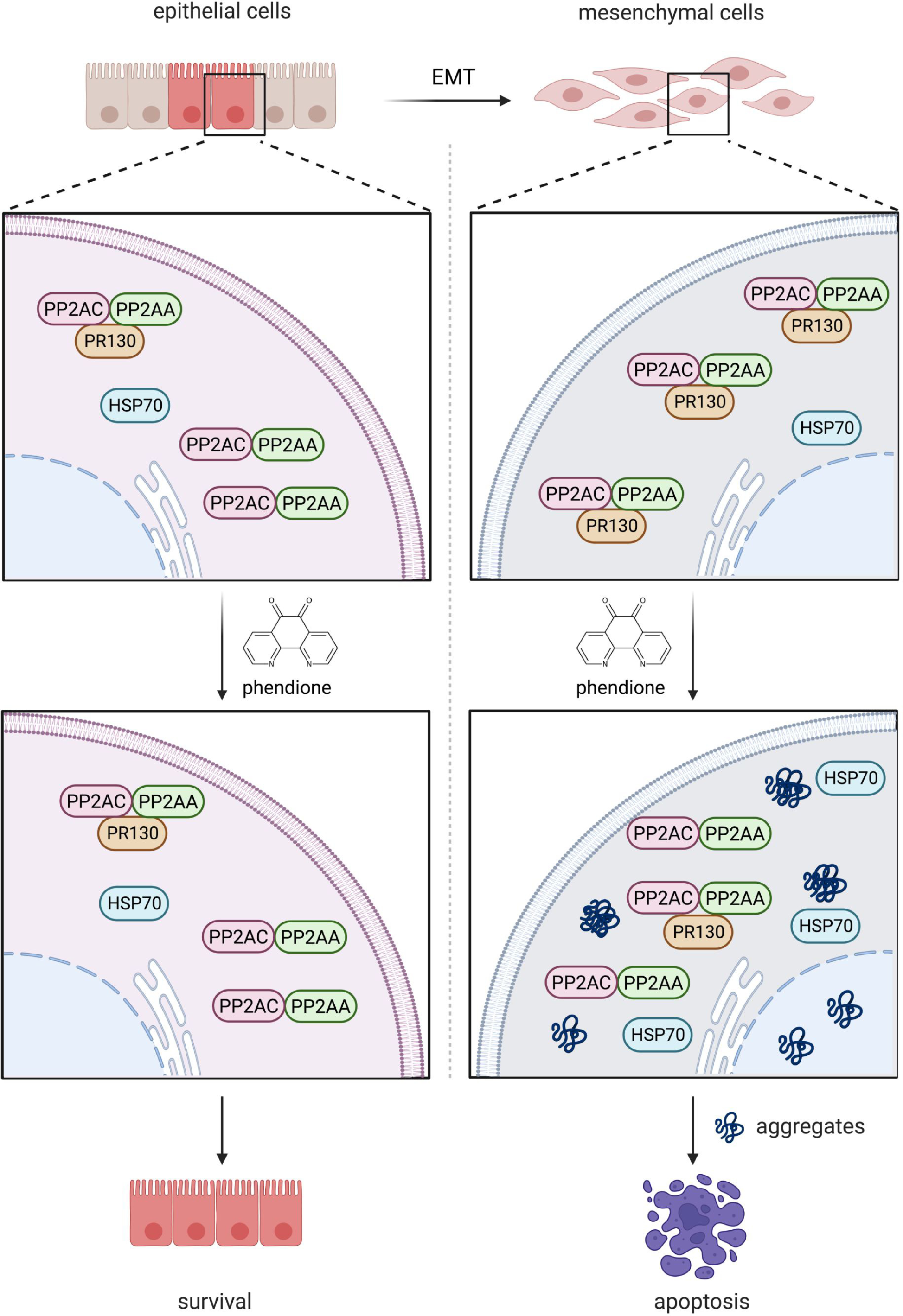

**HIGHLIGHTS:**

➢ The PP2A subunit PR130 is a molecular marker of mesenchymal PDAC cells
➢ The small molecule PP2A inhibitor phendione selectively kills mesenchymal PDAC cells
➢ Phendione decreases PR130 through proteasomes and selectively increases the heat shock protein 70 kDa in mesenchymal PDAC cells
➢ HSP70 promotes cell survival upon inhibition of PP2A
➢ PP2A-PR130 regulates the accumulation of cytotoxic protein aggregates in mesenchymal PDAC cells

## INTRODUCTION

Annual new cancer cases will increase by 21.4% in Europe in the next decade.^1^ Within the malignant neoplasms, pancreatic ductal adenocarcinoma (PDAC) remains a particularly dismal disease. The incidence and mortality of PDAC are increasing and the disease is expected to become the second leading cause of cancer-related deaths, with a current 5-year survival rate of only 12%.^2,3^ In 80-85% of patients with locally advanced or disseminated disease, the DNA-damaging chemotherapeutics folinic acid, fluorouracil, irinotecan and oxaliplatin or nab-paclitaxel and gemcitabine are standards of care.^4^ Such therapies result in overall response rates of only 20-30% and is associated with severe adverse events impacting the quality of life.^4^ Thus, novel therapies to treat PDAC are urgently needed.

To improve the prognosis of PDAC through concepts of precision oncology, it is necessary to characterize PDAC cells better, to develop novel mechanism-based therapies, and to define reliable molecular markers for drug responsiveness. PDAC subtypes with diverse clinical responses to therapies and a differential impact on the survival of patients were identified. These are the squamous/basal-like/quasi-mesenchymal, classic/progenitor, immunogenic, and aberrantly differentiated endocrine exocrine subtypes. The recurrently identified and validated mesenchymal subtype (squamous/basal-like) ties in with the worst patient prognosis. This subtype is associated with early cancer cell dissemination and chemoresistance.^5,6^

Protein kinases catalyze the transfer of ɣ-phosphate groups from ATP molecules to serine, threonine, or tyrosine residues. Kinases are dysregulated in most if not all cancers. This has spurred the development of numerous kinase inhibitors.^7–9^ Depending on the tumor type, these agents produce therapeutic benefits. Phosphatases remove phosphate groups hydrolytically from many physiologically important proteins. This makes phosphatases attractive enzymes for the development of targeted therapies. Their therapeutic potential has yet not been fully exploited.^10^

The protein phosphatase 2A (PP2A) is a major source of cellular serine/threonine phosphatase activity.^11^ Trimeric PP2A complexes contain the scaffolding A subunit (PP2A-A), the catalytically active C subunit (PP2A-C), and one of at least 17 regulatory B-type subunits. B-type subunits regulate the intracellular localization of PP2A and bind individual proteins to confer substrate specificity to PP2A. PP2A is well-known to control cell death by apoptosis, limited cell digestion by autophagy, cell cycle progression, cell proliferation, aging, and differentiation, as well as DNA repair.^11–13^ Far less is known about the biological functions of individual PP2A B-type subunits.^10,11^ This applies to an impact of PP2A on the delicate balance between protein synthesis and degradation.^14^ It is unclear if the accumulation of cytotoxic protein aggregates upon PP2A inhibition, which is seen in neurological diseases and causes the formation of proteinaceous aggregates,^15,16^ occurs in tumor cells. How this accumulation of misfolded proteins affects them, if this is therapeutically exploitable, and if certain PP2A B-type subunits control protein aggregate formation awaits further experimental validation. The clearance of aggregates involves protein degradation by the ubiquitin-proteasome-system,^15^ autophagy,^16^ and the family of heat shock proteins (HSPs) ensuring proper protein (re)folding.^17–19^

The notion that PP2A is frequently inactivated in human cancers and studies in mice suggest that PP2A is a tumor suppressor.^20^ However, PP2A can likewise exert pro-oncogenic functions. For instance, in a murine chemical carcinogenesis model for hepatocellular carcinoma, an overexpression of PP2A-C leads to more and larger tumors.^21,22^ High expression levels of certain PP2A subunits are also linked to cell transformation and the aggressiveness of human tumors. These include the B-type subunits PPP2R2D in gastric cancer and PR130 (encoded by the gene *PPP2R3A*) in liver cancer.^22–26^ A recent work analyzed Cancer-Genome-Atlas (TCGA) datasets and surgically removed human PDACs for a link between the *PPP2R3A* mRNA expression and PR130 with patient survival. PR130 was found to be associated with poor prognosis and with tumor recurrence.^27^ The PP2A B-type subunit *PPP2R5A* was also linked to the metastatic spreading of PDAC cells.^28^ Additional work is required to fully understand how PP2A B-type subunits determine carcinogenesis. For example, it is not clear if PR130 is linked to certain tumor subtypes.

Extending the limited data on the roles of PP2A in PDAC are encouraged by a phase 1 clinical trial of the PP2A inhibitor LB-100 (NCT01837667). LB-100 produced a promising partial remission in one PDAC patient.^29^ Further investigations are obviously needed to evaluate if PP2A inhibition is a valid treatment option. Moreover, it is necessary to find biochemical markers of drug effectiveness. This requires increasingly specific PP2A inhibitors and the identification of cancer cell types that require PP2A for their growth and survival.

Here we demonstrate that mesenchymal PDAC cells express higher levels of PR130 than epithelial PDAC cells. To investigate the relevance of this finding and the associated molecular mechanisms, we used the recently identified small molecule PP2A inhibitor 1,10-phenanthroline-5,6-dione (phendione). We tested phendione because it is more effective and specific for PP2A than LB100.^30^ We identify that an upregulation of the chaperone heat shock protein 70 kDa (HSP70) is a druggable escape mechanism of mesenchymal PDAC cells upon inhibition of PP2A. In cells with inhibited PP2A, PR130 regulates apoptosis and the accumulation of cytotoxic protein aggregates.

## RESULTS

### Differential expression of *Ppp2r3a* and PR130 in epithelial and mesenchymal PDAC cells

We previously analyzed the responses of various genetically defined murine PDAC cells to DNA replication stress inducers and epigenetic drugs.^31^ During this study, we perceived that the protein expression of PR130 varied considerably between different PDAC cell lines. This notion and the fact that very little is known about an association of distinct PP2A B-type subunits with certain tumor types and subtypes let us investigate this systematically. We studied the mRNA expression levels of PP2A B-type subunits in 15 mesenchymal and 23 epithelial cell lines. These cells were recently isolated from genetically defined murine PDACs.^32^ We found that mesenchymal PDAC cells have higher expression levels of the *Ppp2r3a* mRNA encoding PR130 than epithelial PDAC cells (per nomenclature rules, the murine mRNA encoding PR130 is noted as *Ppp2r3a* and the human mRNA encoding PR130 is noted as *PPP2R3A*; the encoded murine and human proteins are both termed PR130). This was specifically seen for *Ppp2r3a* and not for other PP2A B-type subunits and the catalytic/regulatory PP2A subunits *Ppp2ca* and *Ppp2r1a* (Fig. 1a).

**Fig. 1.**
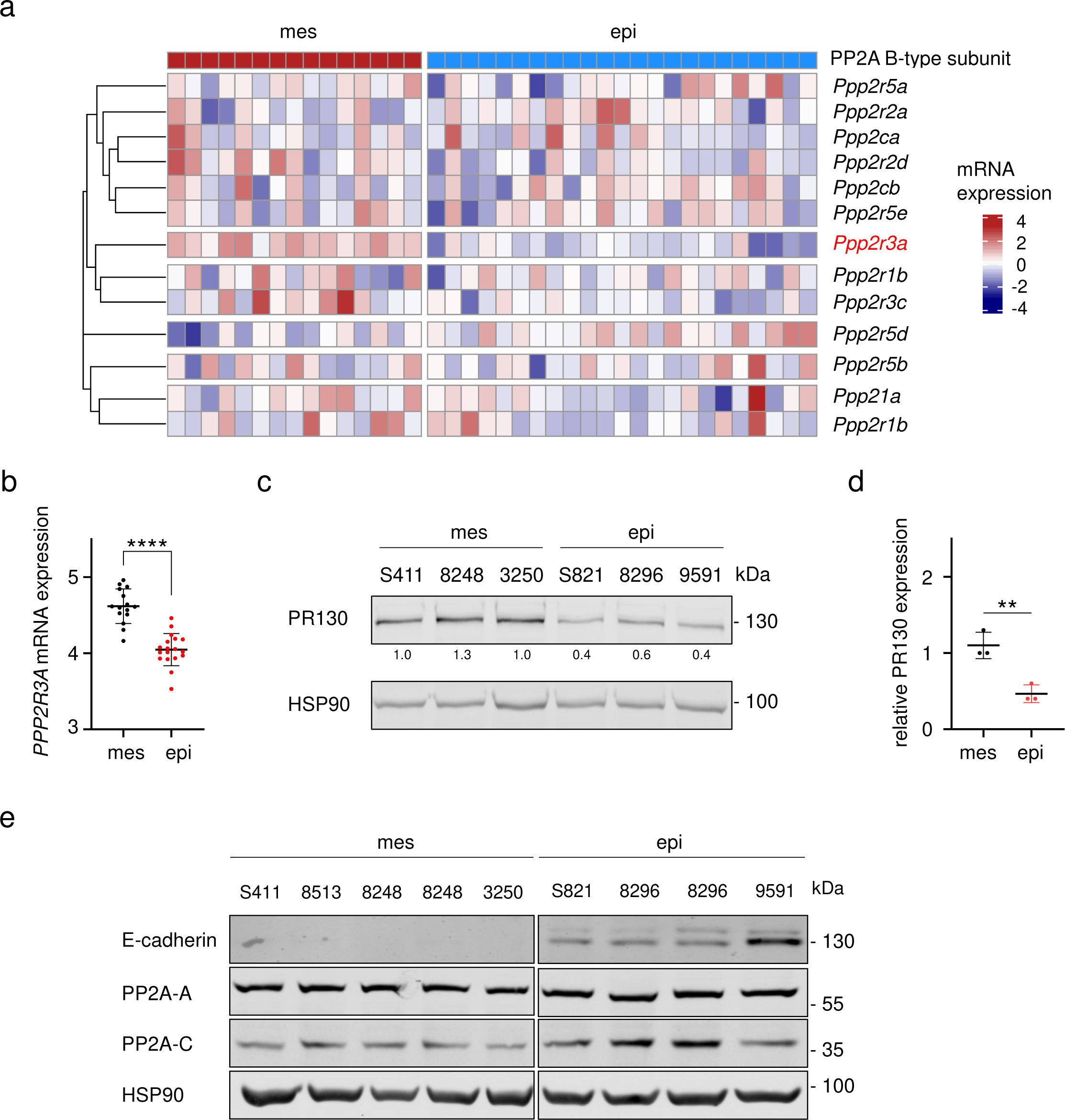
PR130 and *Ppp2r3a* are highly expressed in the mesenchymal PDAC subtype. **a** Heatmap of color-coded mRNA expression data for the depicted PP2A subunits in murine mesenchymal (red, n = 15) and epithelial (blue, n = 23) cell lines by RNA-sequencing. Clustering of rows are based on average distance by pearson correlation (cutree = 6). **b** RNA sequence analyses were done to detect the mRNA expression levels of *Ppp2r3a* in mouse PDAC cell lines; *n* = 15 (mesenchymal) and *n* = 18 (epithelial). Data were statistically analyzed using unpaired t-test, **** *p* < 0.0001). The y-axis is shown as log2-transformed values, normalized to total mRNA contents. **c** Immunoblots show PR130 levels in untreated murine mesenchymal and epithelial PDAC cells. Detection of HSP90 verifies equal sample loading; *n* = 3. **d** Quantification of PR130 detection by immunoblots in mesenchymal and epithelial PDAC cells; unpaired t-test, ** *p* < 0.01). **e** Immunoblot analysis shows the PP2A-A and PP2A-C protein. The detectability of E-cadherin verifies the epithelial origin of the mentioned PDAC cells. 8248 and 8296 cells were blotted twice to also test different passages. HSP90 served as loading control; *n* = 2.

The differences in the levels of *Ppp2r3a* in epithelial and mesenchymal PDAC cells were statistically significant (Fig. 1b). Immunoblot analyses confirmed higher expression levels of PR130 in mesenchymal PDAC cells compared to epithelial murine PDAC cells (Fig. 1c). These differences were also statistically significant (Fig. 1d).

To test if the increased expression of PR130 in mesenchymal is associated with divergent protein levels of the structural PP2A-A and the catalytical PP2A-C subunits, we carried out immunoblots. We found that mesenchymal and epithelial PDAC cells carried similar levels of these major PP2A subunits. We validated the epithelial subtype by detecting the epithelial cell identity marker E-cadherin (Fig. 1e).

These data illustrate that the levels of *Ppp2r3a* and PR130 are a novel molecular marker to discriminate mesenchymal and epithelial PDAC cells.

### Pharmacological inhibition of PP2A induces apoptosis in mesenchymal PDAC cells

The differential expression of PR130 in epithelial and mesenchymal PDAC cells suggests that these subtypes divergently control PP2A-dependent phosphorylation events. We hypothesized that such differences are linked to variable responses of such cells to an inhibition of PP2A. We tested this with the specific PP2A inhibitor phendione.^30^ We treated the mesenchymal cell lines S411 and 8248 and the epithelial cell lines S821 and 8296 with 0.5 µM to 10 µM phendione for 24 h. Microscopical images revealed that ≥ 3 µM phendione caused mesenchymal PDAC cells to round up and detach from the surface of the flasks. Phendione did not alter the morphology of epithelial PDAC cells (Fig. 2a).

**Fig. 2.**
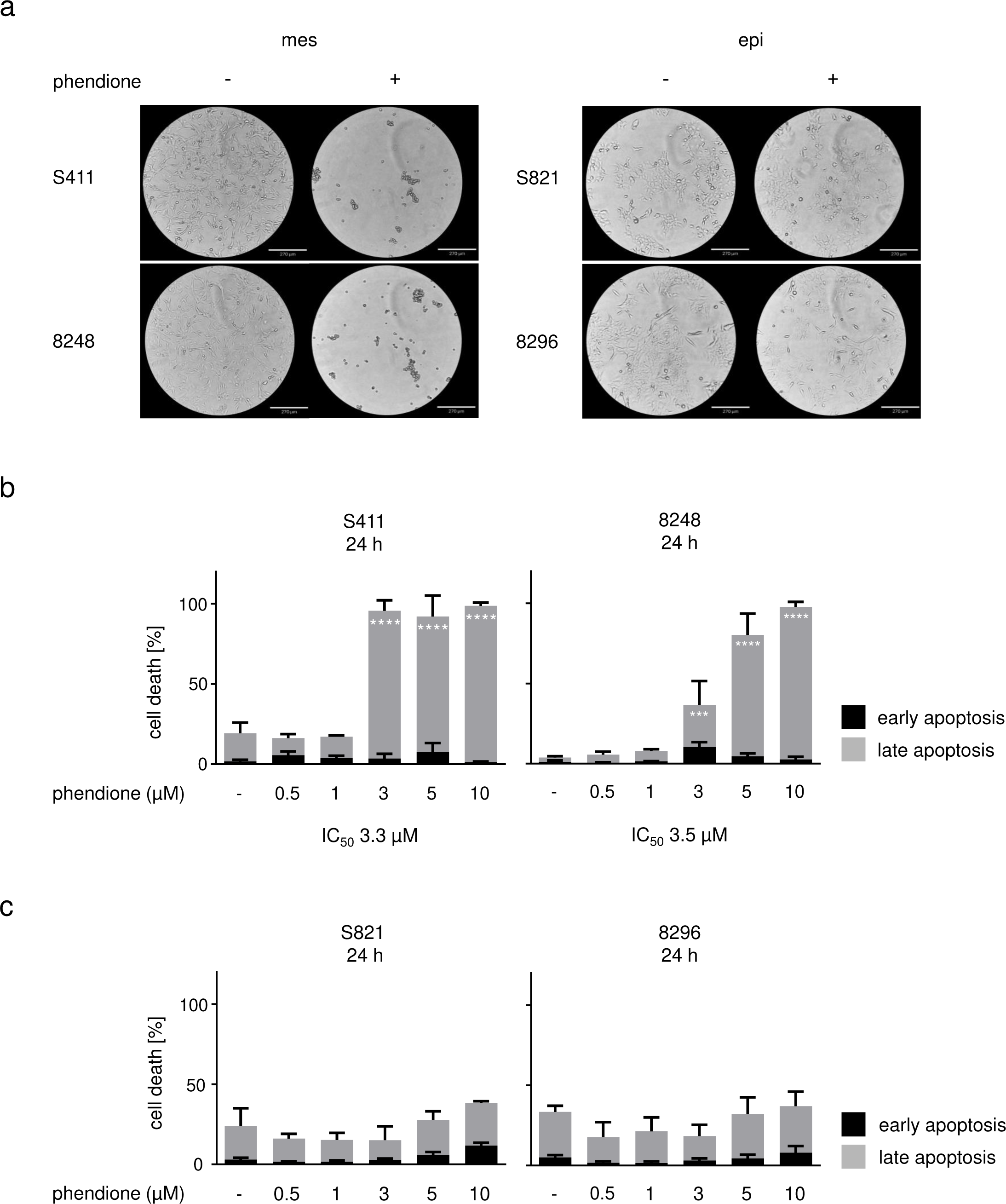
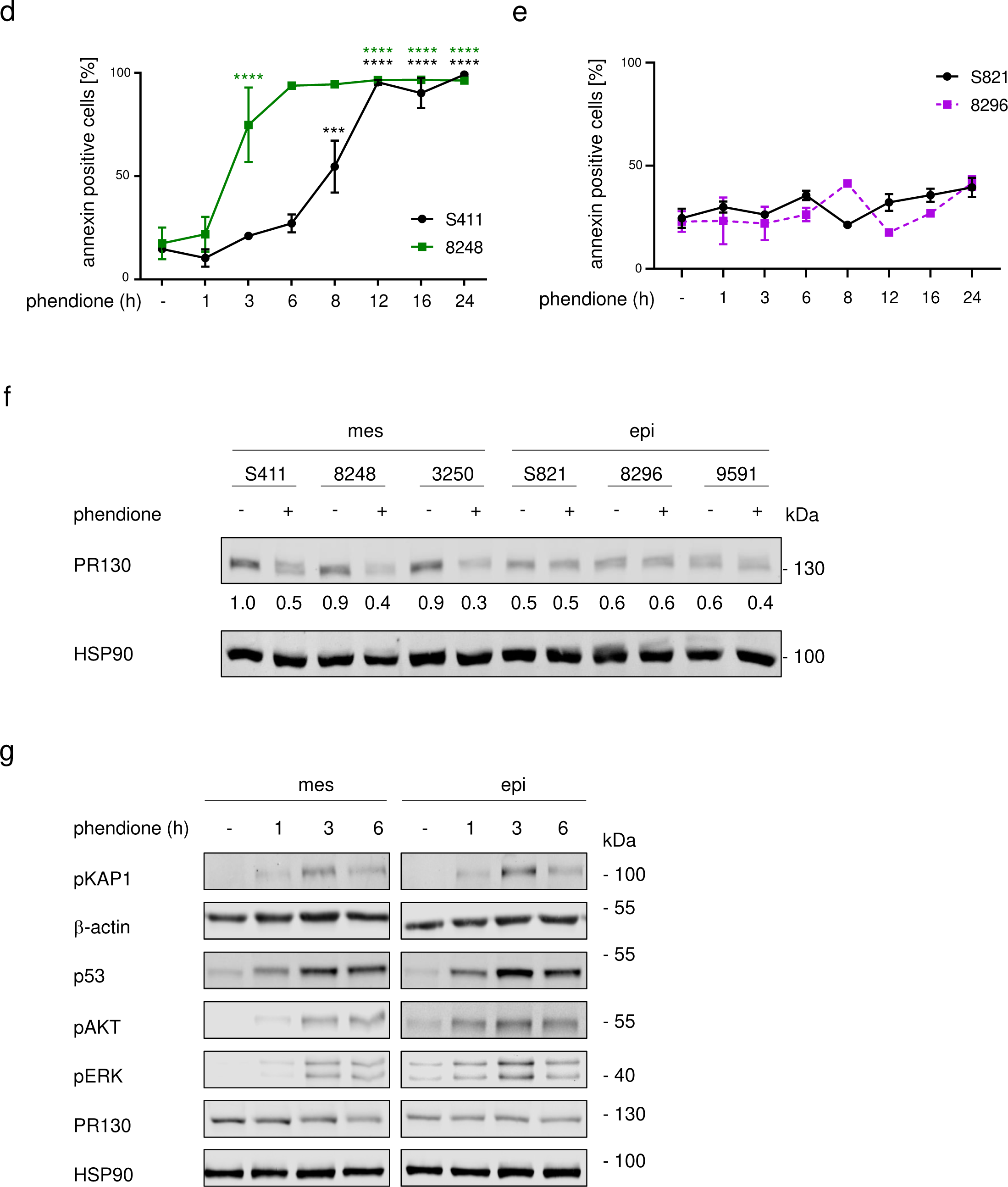
The mesenchymal subtype of PDAC is highly susceptible to the PP2A inhibitor phendione. **a** Morphology of PDAC cells after treatment with 3 µM phendione for 24 h; *n* = 3. **b** Mesenchymal PDAC cells were incubated with concentrations of phendione up to 10 µM for 24 h. Flow cytometry using annexin-V/PI staining was carried out to detect cell vitality and apoptosis; data are shown as mean values ± SD; *n* = 3. **c** Epithelial PDAC cells were treated with 0.5-10 µM phendione for 24 h and apoptosis was determined by flow cytometry for annexin-V/PI; *n* = 3. Data were statistically analyzed using two-way ANOVA (* *p* < 0.05, ** *p* < 0.01, *** *p* < 0.001, **** *p* < 0.0001). **d** Mesenchymal and **e** Epithelial PDAC cells were incubated with 10 µM phendione for up to 24 h. Annexin-positive cells were detected by flow cytometry; *n* = 2. Data were statistically analyzed using unpaired t-test (* *p* < 0.05, ** *p* < 0.01, *** *p* < 0.001, **** *p* < 0.0001). **f** Mesenchymal and epithelial murine PDAC cell lines were treated with 3 µM phendione for 24 h. PR130 was measured by immunoblot. HSP90 served as loading control; *n* = 2. **g** PDAC cells were cultured with 3 µM phendione for 1 h, 3 h, and 6 h. Phosphorylated forms of KAP1, AKT and ERK, as well as total PR130 and p53 were detected by immunoblot. HSP90 and β-actin served as independent loading controls; *n* = 3.

We speculated that such phendione-induced morphological changes of mesenchymal PDAC cells were linked to apoptosis (programed cell death). To test this, we stained the cells with the very sensitive apoptosis marker annexin-V/PI and subjected them to flow cytometry analyses. Annexin-V stains early and late apoptotic cells and PI additionally accumulates in late apoptotic and necrotic cells. Congruent with the results depicted in Fig. 2a, we found that phendione significantly induced apoptosis in mesenchymal PDAC cells (S411 and 8248) (Fig. 2b). After 24 h, 3 µM phendione caused apoptosis in 96% of S411 cell cultures and in 37% of 8248 cell cultures. Annexin/PI-positive 8248 cells increased up to 98% after a 48-h treatment with 3 µM phendione (Supplementary Fig. 1a), indicating that both cell lines show similar time-dependent responses. The IC50 values for phendione-induced apoptosis after 24 h are 3.3 µM for S411 and 3.5 µM for 8248 cells.

Even when they were incubated with 10 µM phendione for up to 48 h, the epithelial S821 and 8296 PDAC cell cultures showed no significant signs of apoptosis (Fig. 2c, Supplementary Fig. 1b). We noted analogous effects in mesenchymal 3250 and epithelial 9591 cells after treatment with phendione for 24-48 h (Supplementary Fig. 1c-f).

To investigate time-dependent apoptosis induction by phendione, we treated mesenchymal and epithelial cell systems with 10 µM phendione for 1 h to 24 h. We used the 3-fold IC50 (Fig. 2b) to detect early apoptotic events. Phendione induced apoptosis in mesenchymal S411 cells significantly after 8 h. The same effect occurred in mesenchymal 8248 cells after only 3 h and in mesenchymal 3250 cells after 6 h, indicating that these cell lines show similar concentration-dependent responses to phendione. Under the same conditions, the epithelial cell lines S821 and 8296 did not undergo apoptosis (Fig. 2d-e). In mesenchymal 3250 PDAC cells, phendione induced apoptosis significantly after 6 h. This effect increased time-dependently (Supplementary Fig. 1g). In epithelial 9591 cells, phendione produced a small increase in early and late apoptosis. This effect vanished over time (Supplementary Fig. 1h).

Analysis of PR130 levels disclosed that phendione decreased PR130 protein levels in mesenchymal PDAC cells but not in epithelial PDAC cells (Fig. 2f). The structural PP2A-A and the catalytical PP2A-C subunits were not affected by phendione in both cell types (Supplementary Fig. 1i).

To control these data, we probed for the anticipated hyperphosphorylation of PP2A target proteins. We found an increased phosphorylation of the ATM target protein KAP1 after a 3-h treatment with phendione in both mesenchymal and epithelial PDAC cells. Moreover, phendione induced an accumulation of the tumor suppressor protein p53, which is stabilized by its phosphorylation. Phendione likewise induced phosphorylated AKT and phosphorylated ERK in both cell lines (Fig. 2g). Thus, we can rule out that the lack of cytotoxic effects of phendione on epithelial PDAC cells is due to a lack of uptake or poor biochemical activity of this drug. We further noted that a 6-h treatment with 3 µM phendione sufficed to reduce PR130 in mesenchymal PDAC cells (Fig. 2g). Together with Fig. 2d and Supplementary Fig. 1g, these data show that phendione-induced apoptosis correlates with a loss of PR130.

A recent study has reported an interplay between PR130 and the tumor suppressor p53 in liver cancer cells.^26^ This may explain why certain PDAC cells are more sensitive to phendione than others. However, like the phendione-induced accumulation of p53 (Fig. 2g), the levels of p53 and its direct target p21 are not associated with either the phendione-sensitive mesenchymal or the phendione-resistant epithelial phenotype (Supplementary Fig. 2a). To further test if p53 is required for phendione-induced apoptosis, we treated the p53-negative mesenchymal murine PDAC cell line W22 with phendione and analyzed apoptosis and proteins of the DNA damage response. Annexin-V/PI measurements showed that 1-10 µM phendione dose-dependently caused apoptosis in W22 cells. ATM and the phosphorylation of its direct targets KAP1 and H2AX were induced, and PR130 was reduced by phendione in such cells. Thus, phendione can trigger pro-apoptotic effects in p53-negative mesenchymal PDAC cells (Supplementary Fig. 2b-c).

These results demonstrate that phendione induces apoptosis in mesenchymal PDAC cells p53-independently and that this process is linked to an attenuation of PR130.

### Phendione produces lethal effects in human PDAC cells

To investigate the role of PP2A/PR130 in human PDAC cells, we used MIA PaCa-2 cells. These cells express epithelial and mesenchymal markers and can serve as a p53 mutant model for the epithelial-mesenchymal transition.^33^ We treated MIA PaCA-2 cells with 0.5-10 µM phendione for 24-48 h, stained them with annexin-V/PI, and subjected them to flow cytometry. This analysis revealed a significant accumulation of cells in early apoptosis. This process started with 3 µM phendione after 24 h and with 1 µM phendione after 48 h. Upon treatment with 10 µM phendione for 48 h, 88% of MIA PaCA-2 cells were in early or late apoptosis. The IC50 value of phendione in MIA PaCA-2 cells is 3.4 µM (Fig. 3a).

**Fig. 3.**
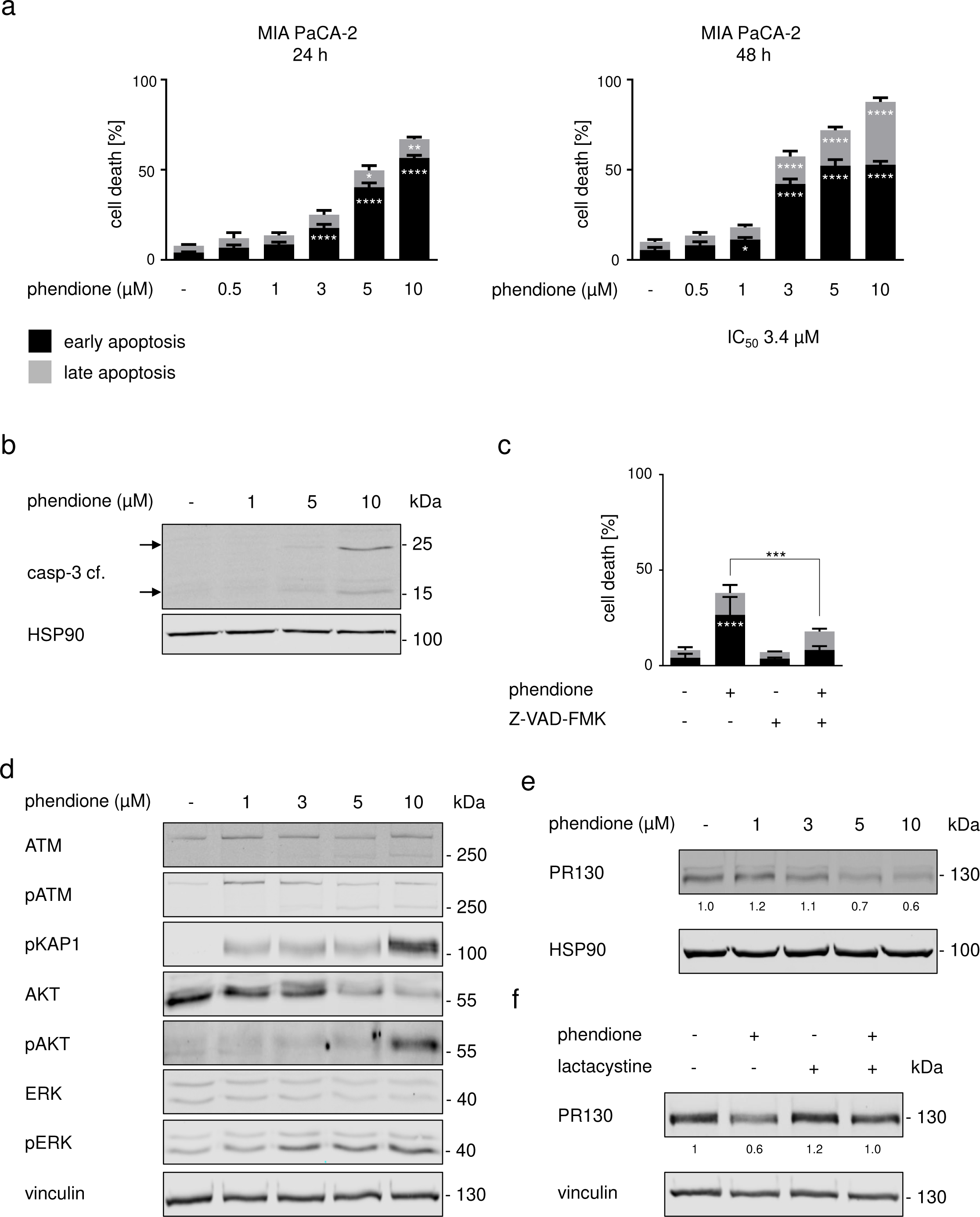
Phendione induces apoptosis in human PDAC cells. **a** The human PDAC cell line MIA PaCA-2 was incubated with up to 10 µM phendione for 24 h and 48 h. Apoptosis was determined by flow cytometry for annexin-V/PI (*n* = 3). Data were statistically analyzed using two-way ANOVA (* *p* < 0.05, ** *p* < 0.01, *** *p* < 0.001, **** *p* < 0.0001). **b** MIA PaCA-2 were treated with increasing doses of phendione up to 10 µM and collected after 24 h. The total and cleaved forms of the apoptosis marker caspase-3 were detected by immunoblot. HSP90 is the loading control; *n* = 2. **c** Apoptosis was measured by flow cytometry of MIA PaCa-2 cells that were exposed for 24 h to 3 µM phendione and 50 µM of the caspase inhibitor Z-VAD-FMK (*n* = 3). Data were statistically analyzed using two-way ANOVA (* *p* < 0.05, ** *p* < 0.01, *** *p* < 0.001, **** *p* < 0.0001). **d** MIA PaCA-2 cells were cultured with 1 µM, 3 µM, 5 µM, and 10 µM phendione for 24 h. Immunoblot was performed for ATM, p-ATM, p-KAP1, AKT, p-AKT, ERK and p-ERK. Vinculin is used as a loading control; *n* = 3. **e** MIA PaCA-2 cells were stimulated with increasing concentrations of phendione (1-10 µM). Immunoblot analyses of whole cell lysates were performed to detect PR130. HSP90 served as loading control; *n* = 3. Quantification was done by normalizing the PR130 signals to those of vinculin. **f** Immunoblot of MIA PaCA-2 cells, which were stimulated with phendione and the proteasomal inhibitor lactacystin was performed to detect PR130. Quantification was done by normalizing the PR130 signals to those of vinculin which was used as loading control; *n = 3*.

To confirm apoptosis induction, we assessed the well-established apoptotic marker cleaved caspase-3 by immunoblot and we used the anti-apoptotic caspase inhibitor Z-VAD-FMK. After 24 h, 5 µM and 10 µM phendione evoked the cleavage of caspase-3. Z-VAD-FMK significantly, though not completely, attenuated the pro-apoptotic effects of phendione in MIA PaCA-2 cells (Fig. 3b-c).

We further verified on-target activities of 1-10 µM phendione in MIA PaCa-2 cells as accumulation of the phosphorylated forms of ATM, KAP1, AKT after 24 h. Moreover, we noted an induction of full-length and the advent of shorter isoforms of ATM and phosphorylated ATM, and that phendione attenuated the total levels of AKT and ERK (Fig. 3d).

Among murine PDAC cell types, PR130 is associated with the highly aggressive mesenchymal phenotype (Fig. 1a-d). PR130 was found to be associated with a shorter survival of PDAC patients.^27^ We investigated if phendione decreased PR130 in human PDAC cells like it did in murine cells (Fig. 2f). Phendione decreased PR130 in MIA PaCa-2 cells dose-dependently (Fig. 3e). Next, we analyzed by which mechanism PR130 is degraded. We incubated MIA PaCA-2 cells with the specific proteasomal inhibitor lactacystin or the autophagy inhibitor chloroquine. Treatment with lactacystin attenuated the phendione-induced reduction of PR130 (Fig. 3f). In contrast, PR130 levels were reduced by phendione irrespective of chloroquine (Supplementary Fig. 3a).

These results illustrate that phendione triggers apoptosis and a proteasomal degradation of PR130.

Phendione induces HSP70 and protein aggregates selectively in mesenchymal PDAC cells To define the molecular mechanisms underlying the differential responses of mesenchymal and epithelial PDAC cells to phendione, we carried out proteomics. We labeled the mesenchymal S411 cells and the epithelial 8296 cells with stable amino acid isotopes and treated them with 2 µM phendione for 24 h (such treatment induces 30% apoptosis in S411 cells after 24 h, see below, Fig. 4f). This analysis revealed a uniquely strong induction of the HSP70 isoform HSP70-1/HSP72 (encoded by the *HSPA1A* gene), and to a lesser extent of its 90% homologous isoform HSP70-1L (encoded by the *HSPA1L* gene) in S411 cells upon treatment with phendione. In 8296 cells, phendione slight induced HSP70-1 and not HSP70-1L (Fig. 4a). Immunoblots with an antibody that recognizes HSP70, but not the related HSC70, confirmed these observations in three mesenchymal and three epithelial murine PDAC cell lines that were treated with 3 µM phendione for 24 h. All tested mesenchymal PDAC cells accumulated HSP70. Epithelial PDAC cells did not accumulate HSP70 upon phendione treatment (Fig. 4b). We further noted that a 6 h treatment with 3 µM phendione sufficed to induce HSP70 in mesenchymal PDAC cells (Fig. 4c).

**Fig. 4.**
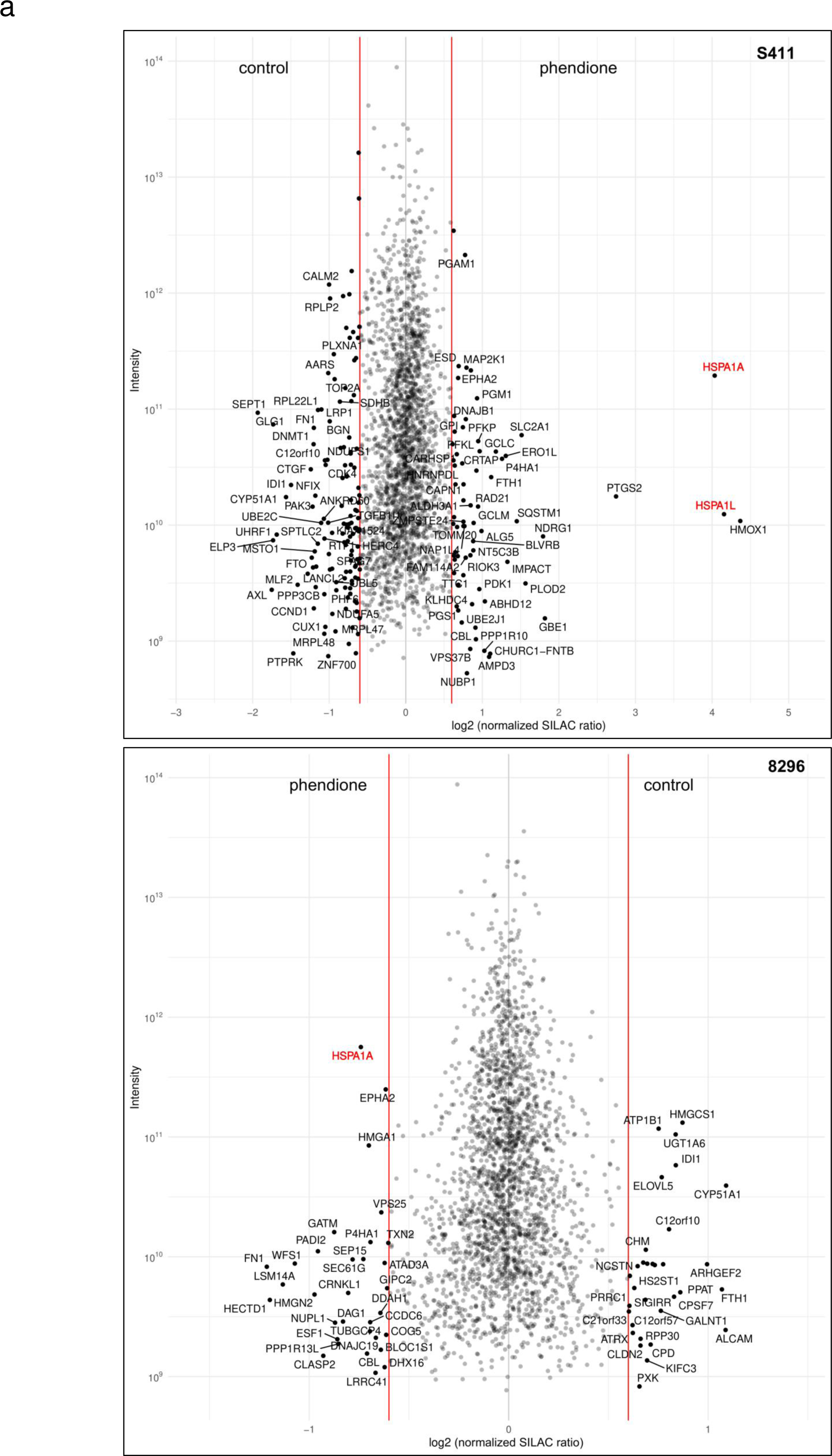
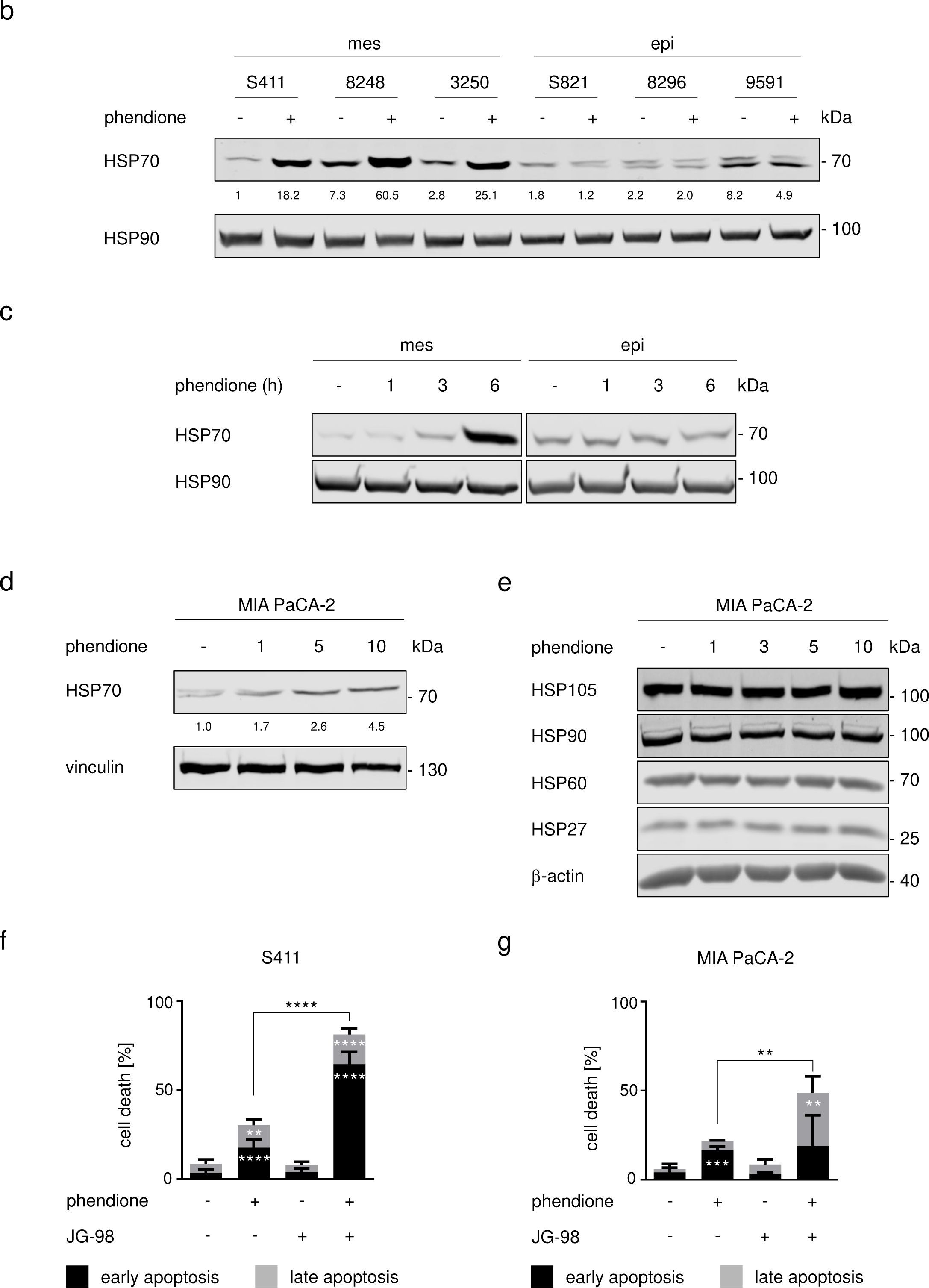
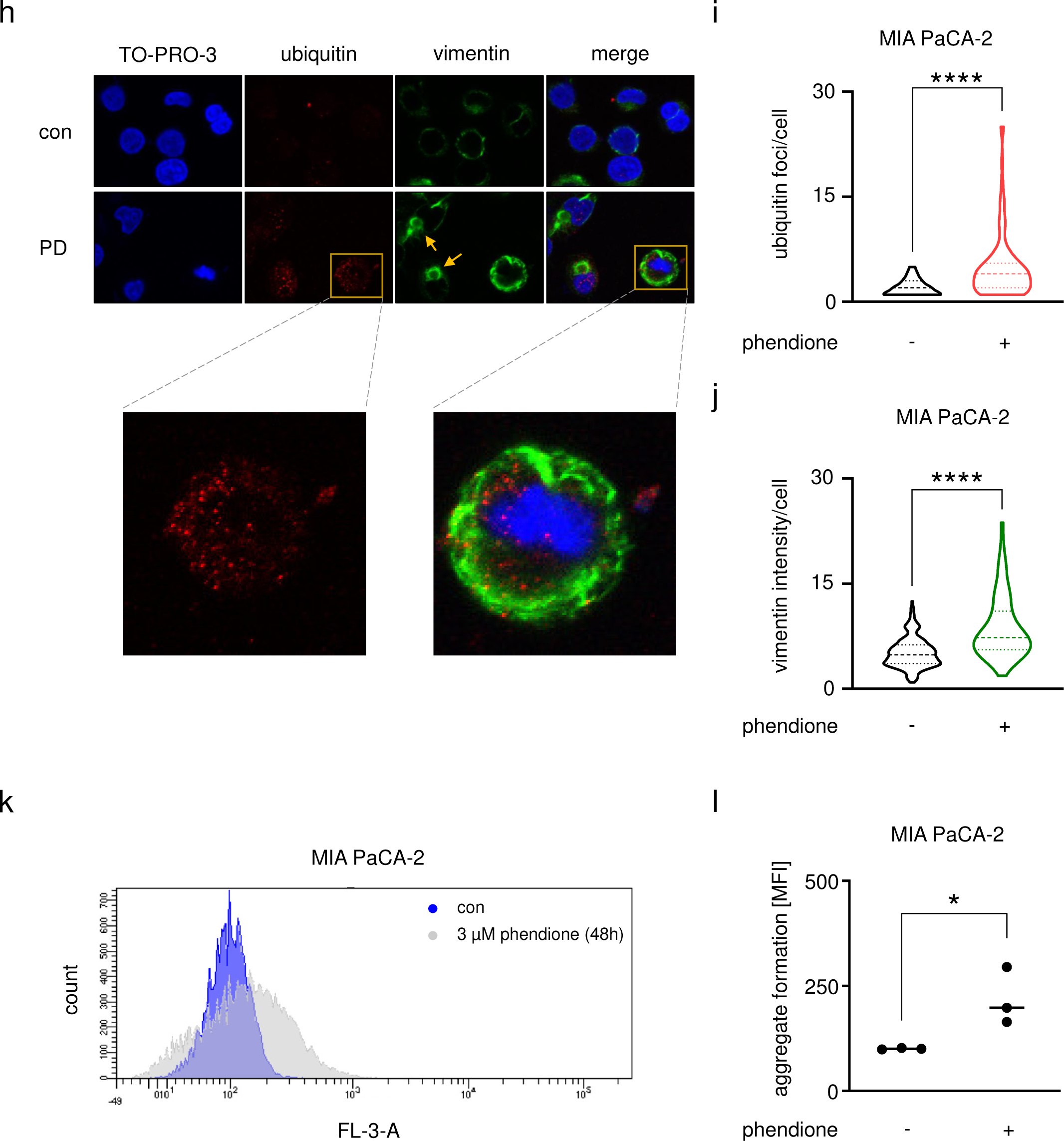
Phendione treatment of PDAC cells leads to an induction of the heat shock protein HSP70. **a** The phendione-sensitive cell line S411 and the phendione-resistant cell line 8296 were labeled with stable amino acid isotopes in cell culture media. The global protein expression profiles of the PDAC cell lines that were incubated with 2 µM phendione for 24 h were evaluated by proteomics (L = light medium; H = heavy medium; PD = phendione). **b** Murine PDAC cell lines were treated with 3 µM phendione for 24 h. Whole cell extracts were blotted for HSP70. HSP90 served as loading control; *n* = 3. **c** Murine mesenchymal S411 cells and epithelial 8296 PDAC cells were cultured with 3 µM phendione for 1 h, 3 h, and 6 h. Immunoblot was done to detect HSP70. HSP90 served as loading control; *n* = 3. **d** Human MIA PaCA-2 PDAC cells were cultured with 1 µM, 5 µM, and 10 µM phendione for 24 h. Immunoblot detected HSP70 and vinculin as loading control; *n* = 3. **e** Immunoblot was done to detect HSP27, HSP60 and HSP105 in MIA PaCA-2 PDAC cells that were treated with 1 µM, 3 µM, 5 µM, and 10 µM phendione for 24 h. HSP90 and β-actin are used as independent loading controls; *n* = 3. **f** Mesenchymal murine PDAC cells (S411) and **g** human PDAC cells (MIA PaCA-2) were incubated with 2 µM phendione and 0.5 µM of the HSP70 inhibitor JG-98 for 24 h. Apoptosis was measured by flow cytometry using annexin-V/PI staining; *n* = 3. Data were statistically analyzed using two-way ANOVA (* *p* < 0.05, ** *p* < 0.01, *** *p* < 0.001, **** *p* < 0.0001). **h** MIA PaCA-2 cells were left untreated (con) or were treated with 3 µM phendione for 24 h (PD), fixed, and incubated with anti-vimentin or anti-ubiquitin antibodies. Alexa Fluor-488 (green, vimentin) and Cy3 (red, ubiquitin) coupled antibodies were used for detection and TO-PRO3 was used to visualize the nuclei. Representative images are shown; *n* = 2. Analysis of vimentin/ubiquitin foci per cell was done using ImageJ software. Data are shown as mean ± SD (one-way ANOVA * *p* < 0.05, ** *p* < 0.01, *** *p* < 0.001, **** *p* < 0.0001; scale bar, 10 µm). **i** Quantification of ubiquitin foci in phendione treated MIA PaCA-2 cells. Data were collected by counting 100 cells in each of 2 independent runs. **j** Quantification of vimentin intensity in phendione-treated MIA PaCA-2 cells. Data were collected by measuring 100 cells in each of 2 independent runs. **k** MIA PaCA-2 cells were seeded and treated with phendione for 48 h; con, untreated. The cells were collected and analyzed using the aggresome detection kit. Data were collected by flow cytometry; *n* = 3. **l** Quantification of data from (k). Data were statistically analyzed using one-way ANOVA (* *p* < 0.05, ** *p* < 0.01, *** *p* < 0.001, **** *p* < 0.0001).

To test whether phendione induces HSP70 in human cells, we treated MIA PaCA-2 cells with 1-10 µM phendione for 24 h. Although to a lesser extent than in murine PDAC cells, MIA PaCa-2 cells accumulated HSP70 upon phendione treatment (Fig. 4d). When we screened lysates from phendione-treated MIA PaCA-2 for other HSPs by immunoblot, we noted that phendione did not modulate HSP105, HSP60, and HSP27 (Fig. 4e). These data suggest a specific control of HSP70 in both murine and human PDAC cells. This accumulation of HSP70 correlates with the reduction of PR130 (Figs. 2g, 3e).

To analyze the biological role of HSP70, we treated PDAC cells with phendione in combination with the HSP70 inhibitor JG-98.^34^. Flow cytometry assessing apoptosis showed that JG-98 alone did not affect cell viability, excluding off-target toxicity of this agent. The inhibition of both PP2A and HSP70 synergistically induced apoptosis in S411 and MIA PaCA-2 cell cultures. A single treatment with 2 µM phendione caused 30% apoptosis in S411 cells and this increased to 81% in the combination treatment. In MIA PaCA-2 cell populations, 22% underwent apoptosis in the presence of 2 µM phendione. This number increased to 49% when 0.5 µM JG-98 was added (Fig. 4f-g). S411 cells are more susceptible than MIA PaCa-2 cells to phendione±JG-98, and this parallels the higher accumulation of HSP70 (Fig. 4b-d). When we treated epithelial 8296 cells with 2 µM phendione and 0.5 µM JG-98, we found that this treatment did not induce apoptosis (Supplementary Fig. 4a). This finding correlates with the notion that phendione does not induce HSP70 in such cells (Fig. 4b).

HSP70 is a molecular chaperone of the protein quality control network (proteostasis). It attaches to its protein substrates and stabilizes them against denaturation or aggregation until conditions improve. Moreover, the HSP70-associated system can direct proteins for ubiquitin-mediated proteasomal degradation and it can resolve cytotoxic protein aggregates that develop upon compromised protein synthesis and stability.^18,19^ To test the formation of such aggregates, we analyzed vimentin microscopically. Vimentin forms cage-like and spherical structures around the cell body when aggregates accumulate. Moreover, we analyzed ubiquitin for its central role in the sorting and labeling of proteins that are to be degraded in the cell.^35,36^ Phendione significantly triggered the formation of ubiquitin foci and vimentin cages in MIA PaCa-2 cells. These surrounded the cell or appeared as perinuclear structures (Fig. 4h-i). The staining intensity of vimentin also increased significantly compared to untreated control cells (Fig. 4j). This was not due to an increased expression of vimentin (Supplementary Fig. 4b).

To quantify the formation of aggregates, we conducted a flow cytometric analysis. We treated MIA PaCA-2 cells with phendione and measured the accumulation of aggregates. Phendione caused aggregate formation significantly in such cells (Fig. 4k-l).

These data demonstrate that HSP70 accumulates in phendione-treated PDAC cells. This is linked to protein aggregate formation.

### Knockdown of PR130 sensitizes PDAC cells to cytotoxic effects of phendione

We hypothesized that the higher expression of PR130 and its phendione-induced degradation in mesenchymal PDAC cells are functionally relevant. To test this, we knocked down PR130 with siRNAs. MIA PaCA-2 cells were transfected with control siRNAs or siRNAs against human *PPP2R3A* and then treated with phendione. Based on Fig. 3e, we chose 3 µM phendione. This concentration induced a partial but not full decrease of PR130. Cell death measurement revealed a significantly higher increase in apoptosis in phendione-treated cells lacking PR130 (Fig. 5a). The elimination of PR130 upon siRNA transfection was verified by immunoblot (Fig. 5b). These data suggest an anti-apoptotic role of PR130. Next, we performed siRNA transfections in murine mesenchymal and epithelial cells. We consistently found that siRNAs against murine PR130 sensitized mesenchymal PDAC cells to pro-apoptotic effects of phendione (Supplementary Fig. 5a). In epithelial PDAC cells, the knockdown of PR130 did not sensitize them to phendione (Supplementary Fig. 5b-c).

**Fig. 5.**
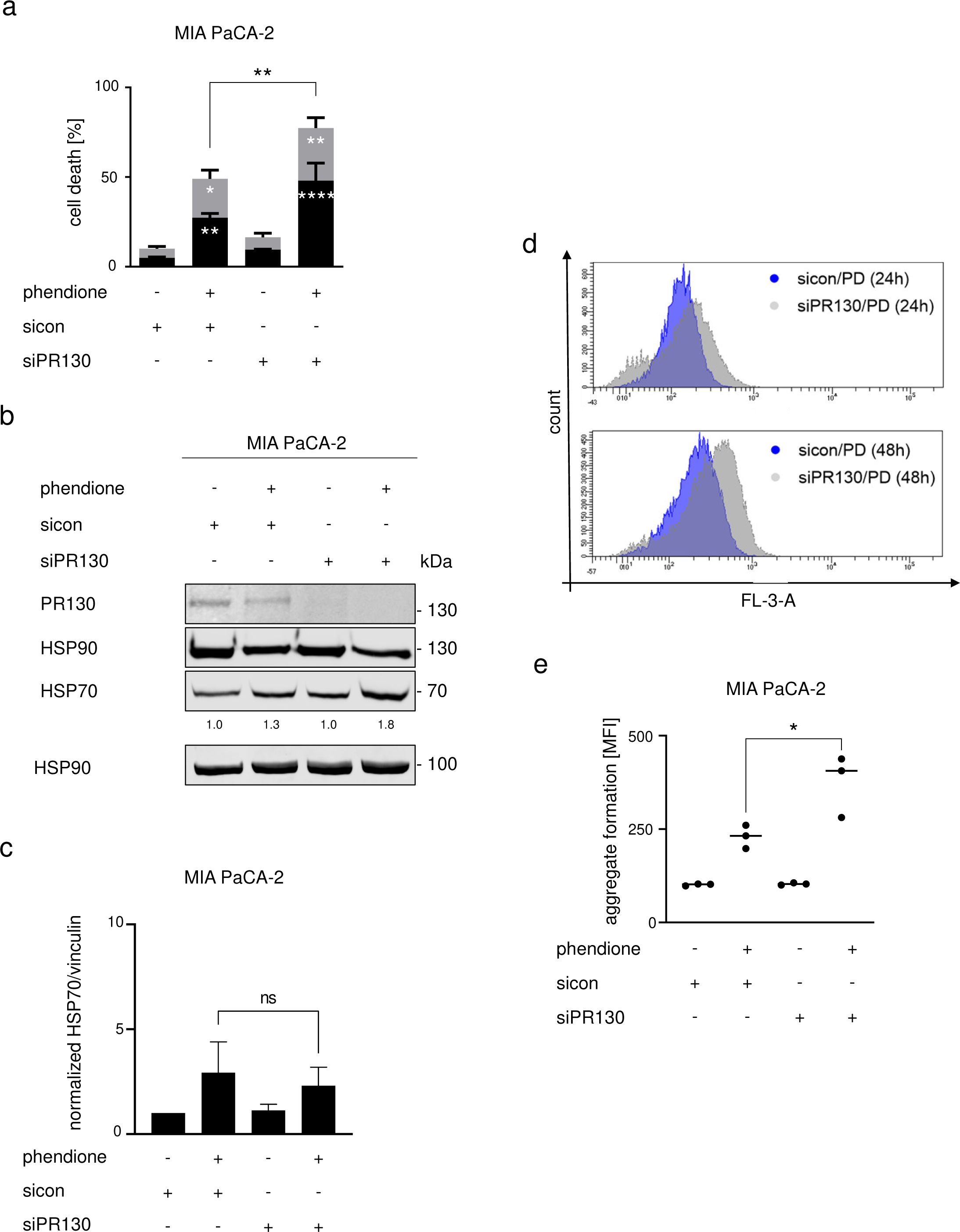
Knockdown of PR130 sensitizes PDAC cells to cytotoxic effects of phendione. **a** MIA PaCA-2 cells were transfected twice within 48 h with 30 pM of the indicated siRNAs (sicon, control siRNAs; siPR130, siRNAs against PR130). After two days, cells were incubated with 3 µM phendione for 48 h. Cell death analysis was done by flow cytometry using annexin-V/PI staining. Data are shown as mean values ± SD; *n* = 3. Data were statistically analyzed using two-way ANOVA (* *p* < 0.05, ** *p* < 0.01, *** *p* < 0.001, **** *p* < 0.0001). **b** siRNA transfection was performed as described in (a). Cells were treated with 3 µM phendione for another 24 h and immunoblot was done to detect PR130 and HSP70. HSP90 served as loading control; *n* = 4. **c** Quantification of HSP70 expression and normalization to the loading control of 4 independent immunoblots from (b). **d** siRNA transfections were performed as described in (a); sicon, control siRNAs; siPR130, siRNAs against PR130. After transfection, MIA PaCA-2 cells were treated with 3 µM phendione (PD) and collected after 24 h and 48 h; con, untreated control cells. Data were analyzed by using the aggresome detection kit and were measured by flow cytometry; *n* = 3. **e** Quantification of data from (c). Data were statistically analyzed using one-way ANOVA (* *p* < 0.05, ** *p* < 0.01, *** *p* < 0.001, **** *p* < 0.0001).

We asked if PR130-depleted mesenchymal PDAC cells mount the HSP70 response upon phendione treatment. Immunoblots showed that untreated and phendione-treated cells lacking PR130 had similar HSP70 levels as control cells (Fig. 5b-c). Having excluded that the elimination of PR130 modulated the accumulation of HSP70, we hypothesized that an elimination of PR130 augmented the phendione-induced aggregate accumulation. To test this, we exposed MIA PaCA-2 cells with and without PR130 to phendione for 48 h and quantified aggregates by flow cytometry. There was a significant difference in the accumulation of aggregates between phendione-treated cells containing PR130 and cells without PR130 (Fig. 5d-e). As control, we performed the same experiment with murine epithelial PDAC cells. These did not build up aggregates and did not induce HSP70 in response to phendione (Supplementary Fig. 5c-d).

This dataset demonstrates that PP2A inhibition induces apoptosis which is linked to the formation of protein aggregates. This process is controlled by PR130.

## DISCUSSION

Reversible phosphorylation is a rapid and straightforward way to modulate biologically relevant protein functions. PP2A represents ∼1% of the cellular protein pool and is a major source of serine/threonine dephosphorylating activity. Specificity of PP2A is largely controlled by B-type subunits that direct catalytically active PP2A complexes to their target proteins.^11,12,20^ This work unravels that the mesenchymal PDAC subtype carries high levels of the PP2A B-type subunit PR130 and has a notable susceptibility to the PP2A inhibitor phendione. Other PP2A B-type subunits and PP2A C and A subunits are though equally expressed in epithelial and mesenchymal PDAC cells. Thus, globally altered PP2A mRNA and protein expression levels in mesenchymal PDAC cells can be ruled out as explanation for the specific vulnerability of mesenchymal PDAC cells towards phendione. This also applies for the major tumor suppressor p53, which is lost or mutated in 60-70% of human pancreatic cancer and mostly in later stages of the disease.^37^ Phendione triggers apoptosis in p53-negative murine PDAC cells and in MIA PaCa-2 cells. These express mutant p53 that is unable to bind p53 consensus DNA sequences. Instead, phendione promotes the accumulation of protein aggregates and decreases PR130 by a proteasome-dependent mechanism. A further reduction of PR130 by RNAi augmented apoptosis induction and the accumulation of protein aggregates. This suggests that PR130-PP2A complexes antagonize cytotoxic protein misfolding and aggregation. Although epithelial PDAC cells respond to phendione biochemically, they are not killed by this agent. They likewise do not accumulate protein aggregates when PP2A is pharmacologically blocked and when PR130 is decreased genetically. We assume that PR130 directs PP2A complexes to proteins that are prone to aggregation in mesenchymal PDAC cells. Such proteins may be linked to migratory phenotypes and metastatic processes.

In neuronal cells, inhibitors of PP2A cause an accumulation of cytotoxic protein aggregates. This can lead to cell death and neuronal disorders, such as in Alzheimer models.^38^ The work presented here reveals that mesenchymal tumor cells accumulate aggregated proteinaceous structures, like neuronal cells do when PP2A functions are impaired. Unlike in neurons, aggregate formation appears to be beneficial in this setting as it can eliminate highly aggressive PDAC cells. In prospective clinical settings with PP2A inhibitors, careful monitoring of neuronal functions might be required.

We conclude that phendione induces the hyperphosphorylation and consequently the misfolding of proteins that are prone for aggregation. The heat shock response is a cytoprotective mechanism against a variety of stressors such as heat shock, oxidative stress, protein misfolding, and heavy metal ions.^17–19^ As a key component of the molecular chaperone system, HSP70 has house-keeping functions. These functions include the folding and assembly of de novo synthesized proteins, protein degradation, refolding of misfolded proteins, and the prevention and removal of protein aggregates. An upregulation of the protein homeostasis regulator HSP70 indicates disturbed proteostasis. Cells can cope with the consequent accumulation of misfolded proteins through their proteasomal degradation.^18,19^ Alterations in vimentin are markers of misfolded proteins and increased proteasome activity.^35,36^ We noted a phendione-induced clustering of vimentin and its increased detectability, which is likely a consequence of such clustering. The occurrence of shorter isoforms of vimentin can be explained by it being a target of caspases-3, -6, and -7 during apoptosis.^39^

The accumulation of protein aggregates indicates that proteasomal degradation is overwhelmed when PP2A is blocked in mesenchymal PDAC cells. This can also be assumed for autophagy, which can eliminate protein aggregates.^16^ The ineffectiveness of this program may likewise be caused by the unexpected lack of co-induction of other HSPs, except for a low increase in DNAJB1/HSP40 that we detected by proteomics. This notion demonstrates that the accumulation of HSP70 upon PP2A inhibition is mechanistically distinct from the well-known selective translation of multiple HSPs when proteotoxic stress is caused by proteasome inhibitors, HSP90 inhibitors, misfolded protein expression, and other proteotoxic stimuli.^17,19^ Why are specifically HSP70-1 and HSP70-1L induced by phendione? Coherent with our data, mesenchymal PDAC sub-populations are very sensitive to inhibitors of protein folding and turnover, including drugs against HSP90.^40^ The serine/threonine phosphatase inhibitor okadaic acid induced the hyperphosphorylation and consequent inactivation of HSP90.^41^ This mechanism might have caused protein aggregate formation in phendione-treated PDAC cells, together with an activation of heat shock factor-1.^42^ This transcription factor selectively activates genes encoding HSPs. Although it is tempting to speculate that such an inhibition of HSP90 induced the accumulation of protein aggregates, our proteome analyses do not show a general decrease of HSP90 target proteins in mesenchymal cells. Moreover, inhibition of HSP90 cannot explain the selective accumulation of HSP70. A recent study shows that LB-100 modulates the RNA splicing machinery.^43^ This may prevent the accumulation of HSPs, except for HSP70 which lacks introns and is not subject to heat shock-repressed translation inhibition.^44^ Our notion that ATM appears in various isoforms in phendione-treated cells may likewise be a result of altered *ATM* RNA splicing that was observed in LB-100 treated colon cancer cells.^43^ Since the levels of total ATM were not altered, caspase-mediated cleavage of ATM,^43^ unlikely explains the occurrence of smaller ATM isoforms.

Beyond its protective role in protein homeostasis, HSP70 provides selective advantages to cancer cells by suppressing multiple cell death pathways, including extrinsic and intrinsic apoptosis and necrosis.^18,19^ We show that a pharmacological inhibition of HSP70 with JG-98 can boost anti-tumor effects of phendione. The mechanism of JG-98 is to block a key allosteric transition of HSP70 that promotes the degradation of some HSP70 target proteins. JG-98 and its analogs are largely selective toward HSP70 family members, demonstrated by pulldowns, overexpression, and point mutation results.^34,45,46^ As a pharmacological inhibition of HSP70 was recently found to block the growth of medulloblastoma cells,^47^ it will be interesting to see how many tumor types are susceptible to a combined inhibition of PP2A plus HSP70. Especially in such a treatment setting, neuronal functions should be monitored carefully.

Phendione-treated PDAC cells with a genetic depletion of PR130 accumulate more protein aggregates than corresponding cells with PR130. In PR130-depleted cells, the induction of HSP70 surprisingly does not increase significantly when PP2A is blocked. Thus, variable accumulation of HSP70 in cells with and without PR130 cannot explain why decreasing PR130 augments apoptosis induction by phendione. We rather interpret this finding as failure of the PR130 knock-down cells to increase HSP70 levels further. It appears that the induction of HSP70 has reached a limit that cannot be increased. Additionally, or alternatively, the loss of PR130 did not allow a further increase in HSP70 despite the occurrence of more aggregates. Obviously, the accumulation of HSP70 cannot prevent aggregates building up and the induction of apoptosis by phendione. Since HSP70 can also promote the poly-ubiquitination and proteasomal degradation of proteins in a complex with the E3 ubiquitin ligase CHIP,^19^ it is possible that this mechanism reduced PR130. Thus, HSP70 may promote protein aggregate removal as well as the loss of PR130 promoting the buildup of such structures. Although these data suggest insufficient molecular mechanisms for aggregates detoxification, the pro-apoptotic effects of the pharmacological inhibition of HSP70 upon PP2A inhibition verify that HSP70 is part of a cellular rescue program.

Previously published data,^27^ and our finding that mesenchymal PDAC cells express more PR130 than epithelial cells suggest that PR130 is a biomarker for reduced overall patient survival. It is currently unclear why mesenchymal murine cells and aggressive human PDACs have higher *Ppp2r3a/PPP2R3A* mRNA and PR130 protein levels. Recent data have demonstrated that cytokines induce the mRNA and protein expression of *PPP2R3A* and PR130 in murine and human cells. These include EGFR signaling in cardiac cells,^48^ and TGF-β1 in pulmonary fibroblast cells.^49^ Such stimuli are associated with tumorigenesis and metastatic tumor cell traits. Thus, they may promote the expression of PR130.

In sum, our findings demonstrate that inhibitors of PP2A and HSP70 kill mesenchymal PDAC cells and that PR130 controls cytotoxic protein aggregate formation. These insights may prospectively offer new approaches and strategies for biomarker identification and a subtype specific, personalized treatment option for PDAC patients.

## MATERIAL AND METHODS

### Drugs and chemicals

1,10-Phenanthroline-5,6-dione, lactacystin, triton X-100, and propidium iodide (PI) were from Sigma-Aldrich Chemie GmbH, Munich, Germany; annexin V-FITC was from Miltenyi Biotec, Bergisch Gladbach, Germany; chloroquine was from Enzo Life Sciences GmbH, Lörrach, Germany; Z-VAD-FMK and the HSP70 inhibitor JG-98 were from Selleck Chemicals, Munich, Germany; dithiothreitol was from PanReac AppliChem, Darmstadt, Germany, formaline (37%) and bovine milk powder were purchased from Carl Roth, Karlsruhe, Germany.

### Cell culture

The isolation and characterization of PDAC cells and details on MIA PaCa-2 cells have been described.^31,32,50^ PDAC cell lines were cultured in high glucose Dulbecco’s Modified Eagle’s medium (D0819, Sigma-Aldrich, Chemie GmbH, Munich, Germany), containing 5-10% fetal calf serum (S0615, Sigma-Aldrich, Chemie GmbH, Munich, Germany) and 1% penicillin/streptomycin (P4333, Thermo Fisher, Frankfurt/Main, Germany). Mesenchymal (S411, 8513, 8248, 3250) and epithelial (8296, S821, 9591) PDAC cells were detached from flasks with 0.5% trypsin-EDTA (15400054, Gibco, Paisley, UK) and re-seeded 1:10 twice per week. The cells were used until passage 12, tested negative for mycoplasma, and MIA PaCa-2 cells were authenticated by DNA fingerprinting (DSMZ, Braunschweig, Germany).

### Protein detection

Immunoblot analyses were carried out as described by us.^31^ Membranes were blocked in 5% milk diluted in TBS-tween-20. Antibodies were diluted in 2% milk/TBS-tween-20. Antibodies were from Cell Signaling, Frankfurt/Main, Germany: PP2A-A (cs-2039), 1:1.000; PP2A-C (cs-2259), 1:1.000; E-cadherin (cs-3795), 1:500; p-AKT (Ser473) (cs-9271S), 1:1.000; p-ERK1/p-ERK2 (Tyr202/Tyr204) (cs-9101), 1:1.000; cleaved caspase-3 (cs-9661), 1:500; ATM (cs-2873), 1:500; ERK1/ERK2 (p44/42) (cs-9102), 1:500; ɣH2AX (Ser139) (cs-9718), 1:1.000; Santa Cruz, Heidelberg, Germany: HSP90 (sc-13119), 1:5.000; p53-DO1 (sc-126), 1:5.000; vinculin (7F9) (sc-73614), 1:5.000; HSP70 (sc-66048), 1:1.000; vimentin (sc-6260), 1:1.000; Novus Biologicals, Wiesbaden, Germany: PR130 (NBP1-87233), 1:500; β-actin (sc-47778), 1:5.000; p-KAP1 (Ser824) (NB100-2350), 1:5.000; Merck, Darmstadt, Germany: ubiquitin (Lys48) (05-1307), 1:500; Abcam, Cambridge, UK: AKT (abcam-32505), 1:1.000; p21 (ab-109199), 1:500 and p-ATM (Ser1981) (ab81292), 1:500; Novocastra Leica Biosystems, Wetzlar, Germany: p53 (NCL-p53-CM5p), 1:500. Secondary antibodies coupled to AlexaFluor-488 (red) for immunofluorescence were from Thermo Fisher (Frankfurt/Main, Germany) and fluorescent secondary antibodies coupled to Cy3 (green) were from Jackson Immuno Research (Cambridgeshire, UK).

### Measurement of cell viability by flow cytometry

Floating and detached cells were collected in FACS tubes and centrifuged at 1.300 rpm for 5 min. The supernatants were discarded, pellets were resuspended in PBS, and centrifuged at 1.300 rpm for 5 min. Afterwards, cells were stained with annexin/V for 20 min at room temperature. Upon adding PI, the samples were measured with a FACS Canto flow cytometer and analyzed with the BD FACSDiva^TM^ Software (BD Biosciences, Heidelberg, Germany). During apoptosis, the plasma membrane undergoes structural changes. This leads to a translocation of phosphatidyl-serine to the extracellular side of the plasma membrane. Annexin-V binds to phosphatidyl-serine and generates a positive detection signal in flow cytometry. Addition of PI allows detection of late apoptotic or necrotic cells, due to the breakdown of the cellular membrane potential and the consequent inability to export PI.

### Genetic knockdown using siRNA

Knockdown of PR130 in murine and human PDAC cells was performed by transfecting 30 pmol of validated siRNA molecules targeting the *PPP2R3A*/*Ppp2r3a* mRNAs (murine: Santa Cruz, sc-108917, 10 µM stock solution; human: Thermo Fisher Scientific, 4392420, 10 µM stock solution) with Lipofectamin® RNAiMAX Reagent (Thermo Fisher Scientific, 13778-075). For control cells corresponding amounts of non-targeting control siRNA-B (Santa Cruz, sc-44230, 10 µM) were used. For each well, 20 µL Lipofectamin® RNAiMAX Reagent, 400 µL Opti-MEM® (Gibco, 31985-070) and siRNA against *PPP2R3A*/*Ppp2r3a* or control siRNA-B were incubated for 10 min at RT. The mixture was added drop by drop to the cells and incubated for 24 h. After 24 h, cells were treated with phendione. Knockdown was verified by immunoblotting.

### Stable isotope labeling with amino acids in cell culture (SILAC) and proteome analyses

PDAC cells were labeled with stable amino acid isotopes in cell culture for 2 weeks. We cultured them in light medium (L-arginine-0 and L-lysine-0) or heavy medium (L-arginine-10 and L-lysine-8). The resulting mass differences could be detected by mass spectrometry. After the period of growing, cells were seeded, and treated with phendione. Floating and detached cells were collected in reaction tubes and centrifuged at 1.300 rpm for 5 min. After washing the pellets with PBS and centrifugation, NuPAGE® LDS Sample Buffer (1x) (Thermo Fisher Scientific, NP0007) with 10% 1 M dithiothreitol was added to the pellets. The samples were incubated for 30 min on ice. Lysates were sonified (10 s), heated (70° C, 10 min), and centrifuged (25 min, 13.000 rpm). Supernatants were transferred to new reaction tubes. Samples were measured by mass spectrometry as described.^51,52^ All proteomics data are deposited to the ProteomeXchange Consortium via the PRIDE partner repository, dataset identifier PXD044854.^53^

### Protein aggregate detection

The aggresome detection kit (ab139486) was from Abcam, Cambridge, UK. Cells were detached by trypsin/EDTA and centrifuged at 1.300 rpm for 5 min. Pellets were washed in 2 mL PBS, centrifuged (5 min, 1.300 rpm), and supernatants were discarded. Pellets were resuspended in 200 µL PBS and added dropwise to 2 mL of a 4% formaldehyde solution (37% formalin in 1x assay buffer). The tubes were mixed carefully to complete the fixation step. After the cell suspensions were incubated for 30 min at RT, a centrifugation step at 800 x g for 15 min was done. The supernatants were removed, pellets resuspended in 2 mL PBS, and centrifuged at 800 x g for 15 min. To permeabilize the cells, 0.5% triton X-100 and 3 mM EDTA (pH = 8) in 1x assay buffer was added by gentle mixing and incubation for 30 min on ice. Afterwards, cells were centrifugated at 800 x g for 15 min and washed with PBS. The cell suspensions were placed in a cell strainer and centrifuged to remove cellular debris. To detect aggregates, 500 µL of freshly prepared aggresome red detection reagent (1:5.000 in 1x assay buffer) was added to the pellets and incubated for 30 min in the dark. Samples were measured with FACS Canto flow cytometer and analyzed with the BD FACSDiva^TM^ Software.

### Immunofluorescence analyses

Cells were seeded and treated on coverslips. After 24 h, the coverslips were washed with PBS once and fixed in an acetone-methanol-solution (3:7) at -20 °C for 8 min. Blocking solution (10% BSA, 0.25% triton X-100 in PBS) was added and incubated for 1 h at RT. Incubation with the primary antibodies was carried out overnight in a wet chamber at 4 °C. Antibodies against ubiquitin (1:100) and vimentin (1:200) were diluted in blocking solution. The next day, the coverslips were washed with PBS (thrice for 5 min) and incubated for 1 h at room temperature with secondary antibodies (1:400, diluted in blocking solution). Cells were washed twice with PBS, once with high salt PBS (0.4 M NaCl in PBS) for 2 min, and once more with PBS. Using a scalpel and tweezers, we carefully placed the coverslips on a slide and added 10 µL Vectashield® (Biozol, Vec-H-1.000) containing TO-PRO^TM^-3 (Thermo Fischer, T3605) in a dilution of 1:100 to stain the nuclei. Images were captured with the confocal microscopy Zeiss Axio Observer.Z1 microscope equipped with a LSM710 laser-scanning unit (Carl Zeiss, Jena, Germany). Analysis was performed with the software ImageJ.

### Statistics

Statistical analyses were carried out using one-way or two-way ANOVA and unpaired t-test from GraphPad Prism Vers.8.3.0 software. Correction of statistical analyses was achieved with Bonferroni multiple comparison; *p* values indicate statistical significance.

## Supporting information

Supplemental figures_bioRxiv

## DATA AVAIBILITY STATEMENT

All data are available in the manuscript and at the ProteomeXchange via the PRIDE database, dataset identifier PXD044854.

## ACKNOLEDGEMENTS

We thank Christina Brachetti and Andrea Piée-Staffa for technical support. Prof. Dr. G. Schneider and Dr. Matthias Wirth were invaluable discussion partners throughout the project.

## AUTHOR CONTRIBUTIONS

A.N. performed immunoblots and flow cytometry analyses. A.K.L. performed immunoblots and flow cytometry analyses. A.M.M. established microscopic analysis for all immunofluorescences. F.B. performed and evaluated the proteomic assay, as well as the design of the volcano blots. J.M. established PDAC cells cell cultures, carried out and quantified RNA sequences data. O.H.K. collected funding, designed the experiments, interpreted the data, supervised the project. A.N. and O.H.K. wrote the manuscript with the help of all authors.

## INSTITUTIONAL REVIEW BOARD STATEMENT

Not applicable.

## INFORMED CONSENT STATEMENT

Not applicable.

