## Supplemental figures_bioRxiv for "The protein phosphatase-2A subunit PR130 is linked to cytotoxic protein aggregate formation in mesenchymal pancreatic ductal adenocarcinoma cells"

**This PDF file includes:**

Figures S1 to S5

**Figure S1**

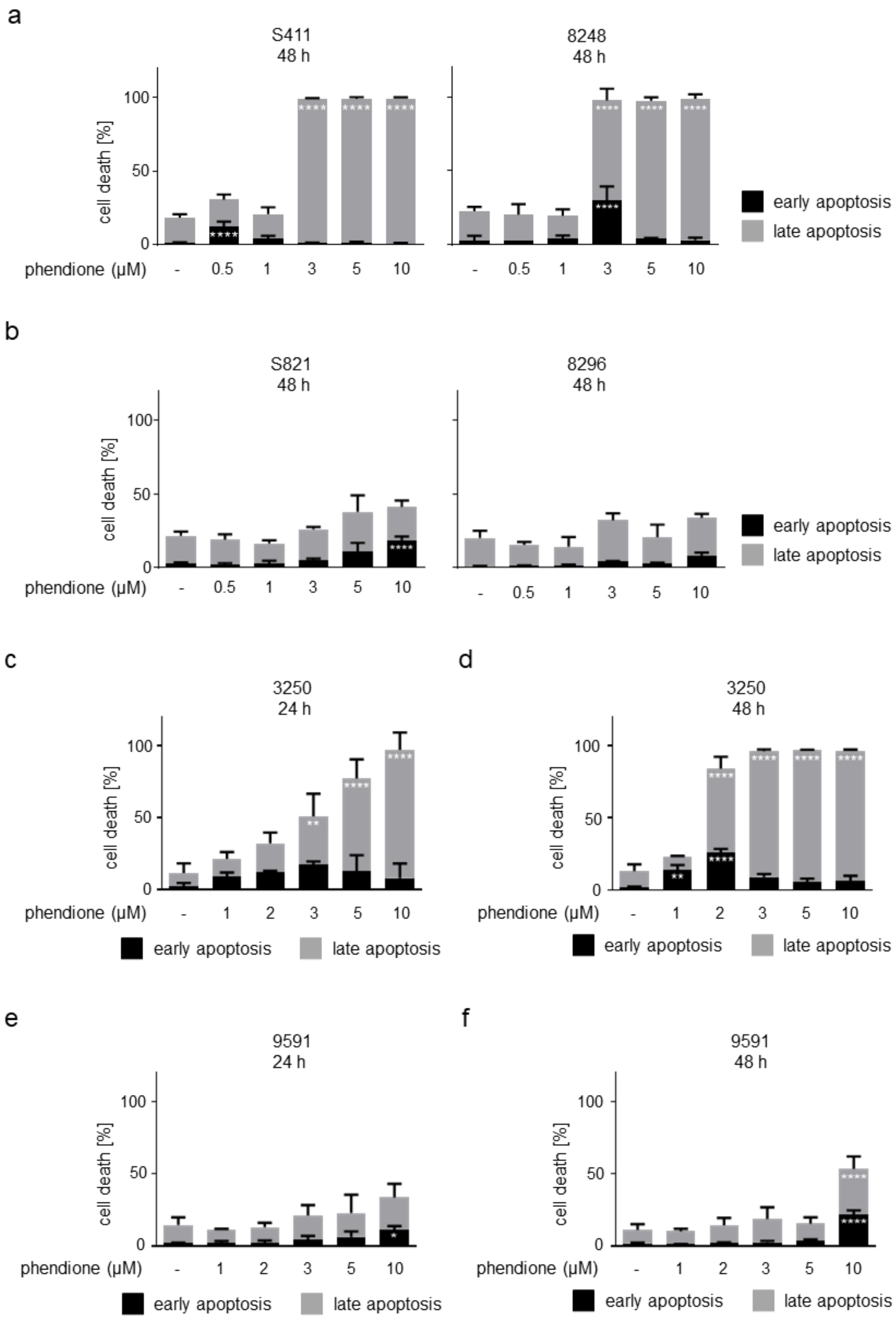
a
S411
48 h
8248
48 h
cell death [%]
100
50
0
phendione (µM)
- 0.5 1 3 5 10
early apoptosis
late apoptosis
b
S821
48 h
8296
48 h
c
3250
24 h
d
3250
48 h
- 1 2 3 5 10
e
9591
24 h
f
9591
48 h

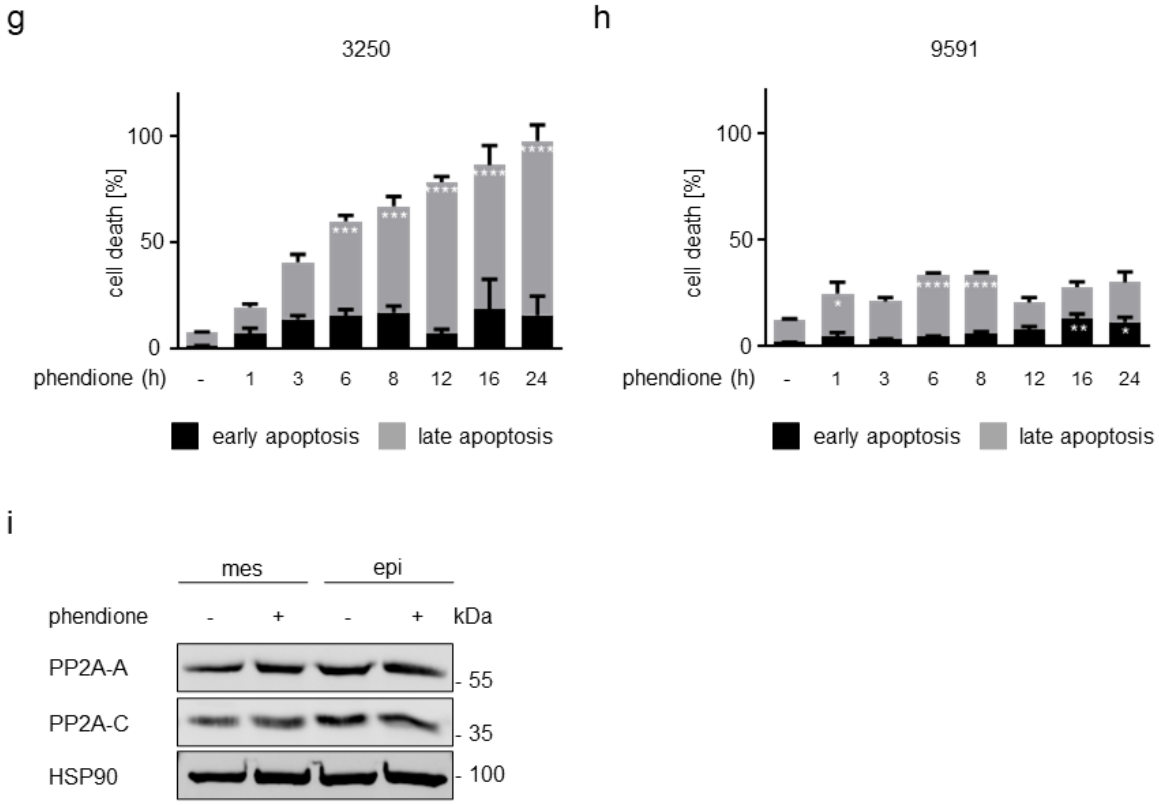


**Supplemental Fig. S1.** Phendione induces apoptosis in mesenchymal PDAC cells in a dose-dependent manner. **a** Apoptosis analyses of mesenchymal PDAC cells after treatment with phendione concentrations up to 10 µM for 48 h. Results were collected by flow cytometry using annexin-V/PI staining and are shown as mean values ± SD; S411: *n* = 3; 8248: *n* = 2. **b** Epithelial PDAC cells were treated with 0.5-10 µM phendione for 48 h and apoptosis was determined by flow cytometry for annexin-V/PI; *n* = 3. **c** Apoptosis analyses were conducted with the murine PDAC cell line 3250 with phendione concentrations up to 10 µM for 24 h. Results were collected by flow cytometry using annexin-V/PI staining and are shown as mean values ± SD; *n* = 3. **d** Apoptosis analyses were carried out with the murine PDAC cell line 3250 with phendione concentrations up to 10 µM for 48 h. Results were collected by flow cytometry using annexin-V/PI staining and are shown as mean values ± SD; *n* = 3. **e** The epithelial PDAC cell line 9591 was treated with 1- 10 µM phendione for 24 h and apoptosis was determined by flow cytometry for annexin-V/PI; *n* = 5. **f** The epithelial PDAC cell line 9591 was treated with 1 – 10 µM phendione for 48 h and apoptosis was determined by flow cytometry for annexin-V/PI; *n* = 4. **g** The mesenchymal PDAC cell line 3250 was stimulated with 10 µM phendione for up to 24 h. Cell death was measured by flow cytometry using annexin-V/PI staining and flow cytometry; $n = 2$. **h** The epithelial PDAC cell line 9591 was incubated with 10 µM phendione for up to 24 h. Apoptosis was measured by annexin-V/PI staining and flow cytometry; $n = 2$. All data (S1 a-h) were statistically analyzed using two-way ANOVA (* $p < 0.05$, ** $p < 0.01$, *** $p < 0.001$, **** $p < 0.0001$). **i** Mesenchymal and epithelial murine PDAC cell lines were treated with 3 µM phendione for 24 h. PP2A subunit A and C were measured by immunoblot. HSP90 served as loading control; $n = 3$.

**Figure S2**

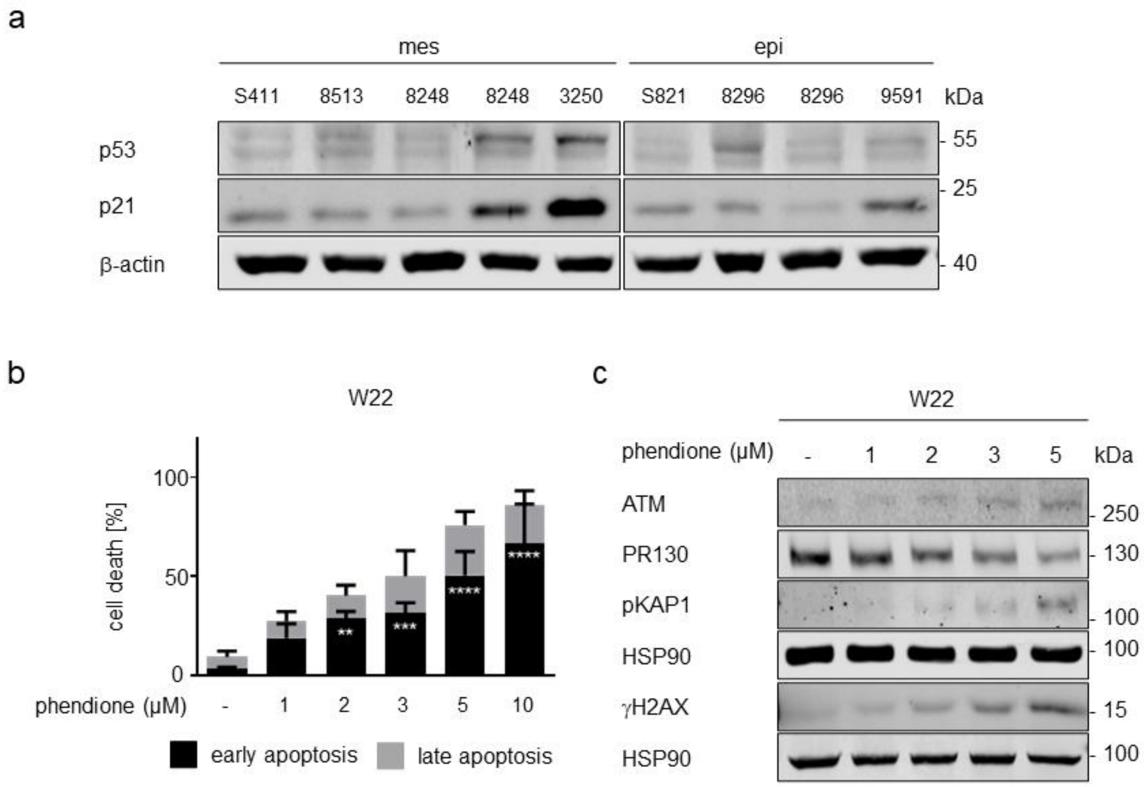


**Supplemental Fig. S2.** Apoptosis induction in phendione-treated cells is p53-independent. **a** Immunoblot analyses show p53, and p21. β-actin serves as loading control; *n* = 2. **b** The p53 null PDAC cell line W22 was treated with 1-10 µM phendione for 24 h. Cell death was determined by flow cytometry; *n* = 3. Data were statistically analyzed using two-way ANOVA (* $p < 0.05$, ** $p < 0.01$, *** $p < 0.001$, **** $p < 0.0001$). **c** Immunoblot analysis was performed with lysates from W22 cells that were incubated with 1-5 µM phendione for 24 h. Immunoblot was done for indicated proteins. HSP90 served as loading control; *n* = 2.

**Figure S3**

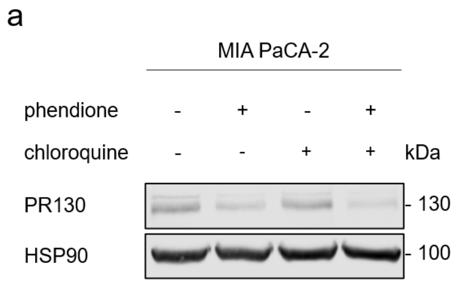


**Supplemental Fig. S3.** Chloroquine is not capable of stabilizing PR130 in phendione-treated cells. **a** Human PDAC MIA PaCA-2 cell line were treated with 3 µM phendione ± 50 µM chloroquine for 24 h. Immunoblot was done to detect PR130 levels. HSP90 served as loading control; *n* = 3.

**Figure S4**

a

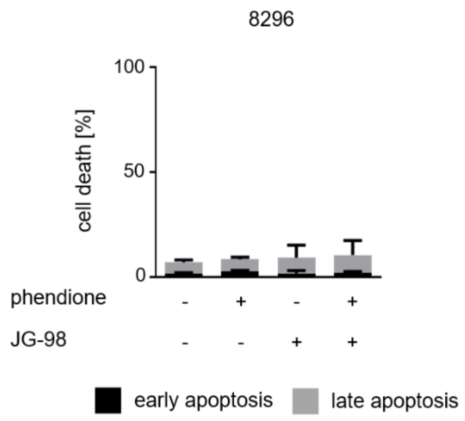


b

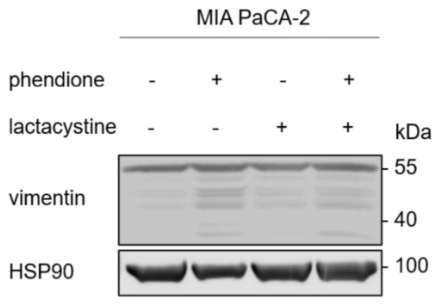


**Supplemental Fig. S4. a** The epithelial murine PDAC cell line 8296 was incubated with 2 µM phendione ± 0.5 µM of the HSP70 inhibitor JG-98 for 24 h. Apoptosis was measured with annexin-V/PI-stained cells and flow cytometry; *n* = 3. Data were statistically analyzed using two-way ANOVA (* $p < 0.05$, ** $p < 0.01$, *** $p < 0.001$, **** $p < 0.0001$). **b** Vimentin status in MIA PaCA-2 cells was analyzed by immunoblot. Cells were seeded, treated with 3 µM phendione ± 50 µM lactacystin and harvested after 24 h. HSP90 served as loading control; *n* = 3.

**Figure S5**

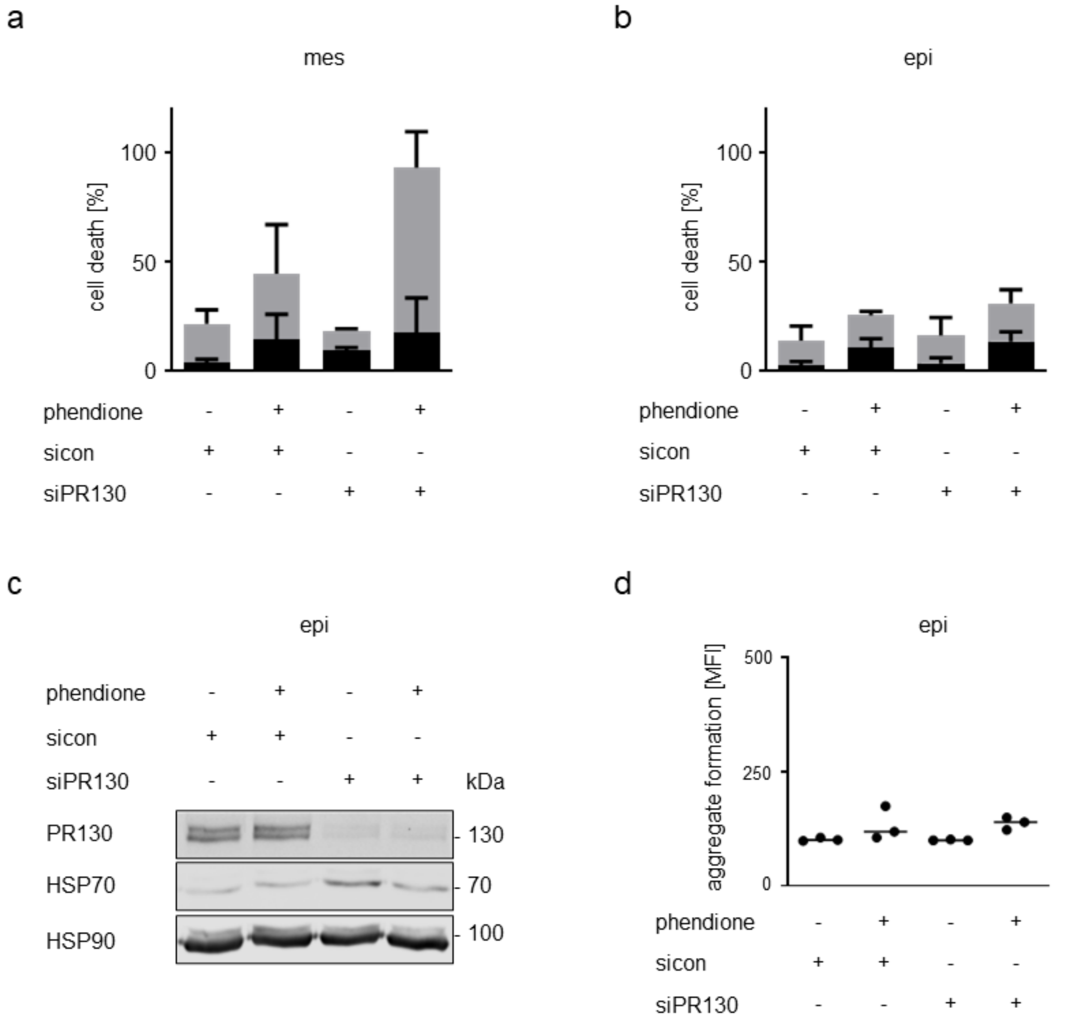


**Supplemental Fig. S5.** Knockdown of PR130 sensitizes mesenchymal but not epithelial murine PDAC cells to toxic effects of phendione. **a** Knockdown of PR130 in mesenchymal PDAC cells was performed as described in the main text. Cells were treated with 3 µM phendione. After 24 h, the cells were harvested and cell death was determined by flow cytometry; $n = 2$. **b** Murine epithelial cells were transfected with siRNAs against PR130 as described in the main text and treated with 3 µM phendione for 24 h. Apoptosis was measured by annexin-V/PI staining and flow cytometry; $n = 2$. **c** The murine epithelial PDAC cell line was transfected and treated as described in (B). Immunoblot was done to detect PR130 and HSP70. HSP90 served as loading control; $n = 3$. **d** 8296 cells were transfected and treated as described above. The accumulation of aggresomes was analyzed using the aggresome detection kit. Data were measured by flow cytometry and statistically analyzed using two-way ANOVA (* $p < 0.05$, ** $p < 0.01$, *** $p < 0.001$, **** $p < 0.0001$); $n = 3$.
